# Synthetic mucus barrier arrays as a nanoparticle formulation screening platform

**DOI:** 10.1101/2023.11.29.569212

**Authors:** Harry Zou, Allison Boboltz, Yahya Cheema, Daniel Song, Gregg A. Duncan

**Author notes:** Correspondence to: Gregg Duncan. These authors contributed equally to this work (Harry Zou, Allison Boboltz).

## Abstract

A mucus gel layer lines the luminal surface of tissues throughout the body to protect them from infectious agents and particulates. As a result, nanoparticle drug delivery systems delivered to these sites may become trapped in mucus and subsequently cleared before they can reach target cells. As such, optimizing the properties of nanoparticle delivery vehicles, such as their surface chemistry and size, is essential to improving their penetration through the mucus barrier. In previous work, we developed a mucin-based hydrogel that has viscoelastic properties like that of native mucus which can be further tailored to mimic specific mucosal tissues and disease states. Using this biomimetic hydrogel system, a 3D-printed array containing synthetic mucus barriers was created that is compatible with a 96-well plate enabling its use as a high-throughput screening platform for nanoparticle drug delivery applications. To validate this system, we evaluated several established design parameters to determine their impact on nanoparticle penetration through synthetic mucus barriers. Consistent with the literature, we found nanoparticles of smaller size and coated with a protective PEG layer more efficiently penetrated through synthetic mucus barriers. In addition, we evaluated a mucolytic (tris (2-carboxyethyl) phosphine, TCEP) for use as a permeation enhancer for mucosal drug delivery. In comparison to N-acetyl cysteine (NAC), we found TCEP significantly improved nanoparticle penetration through a disease-like synthetic mucus barrier. Overall, our results establish a new high-throughput screening approach using synthetic mucus barrier arrays to identify promising nanoparticle formulation strategies for drug delivery to mucosal tissues.

## INTRODUCTION

Mucus is continuously produced to form a protective layer in mucosal tissues throughout our body to prevent irritation, infection, and injury.^1–3^ To eliminate pathogenic and other hazardous materials, mucus physically blocks and/or chemically binds to micro– and nanoscale particles depending on their size and surface chemistry. It has been shown that these barrier functions of mucus can limit the bioavailability of nanoparticle (NP) formulations given orally or administered locally to the eye, nose, lung, and vaginal tract.^4–7^ For example, previous work has demonstrated nanoparticles with hydrophobic properties are strongly adherent to the mucus gel and quickly eliminated from the respiratory and reproductive tract.^8,9^ In contrast, NP formulated with a dense surface coating of polyethylene glycol (PEG) were found to efficiently penetrate the mucus barrier and widely distribute within mucosal tissues. Enhancements in mucus penetration by PEGylated NP can be largely attributed to their near-neutral charge and hydrophilic surfaces which avoid adhesive interactions with the net-negatively charged and hydrophobic regions of mucin glycoproteins.^10,11^

In addition to PEGylation, several alternative strategies have been explored to facilitate NP passage through the mucus barrier such as peptide and zwitterionic polymer coatings as well as the co-administration of NP with mucus-degrading agents (mucolytics).^12–16^ Moreover, the optimal dimensions of NP formulations (e.g. effective diameter, shape) can also vary depending on the target tissue. For example, the characteristic pore size of the mucus barrier can vary from as low as 20 nm in the adherent mucus layer in the gastrointestinal (GI) tract to up to 500 nm in the cervicovaginal tract.^17,18^ In comparison to traditionally spherical NP, prior work has also demonstrated enhanced penetration of rod-shaped NP through mucus in the GI tract.^19,20^ It is also important to note mucus barrier properties can be significantly altered as a function of disease which may lead to improved or limited NP penetration to the underlying tissue.^21–23^ This prior research highlights the numerous concepts and approaches that one may consider in the design of NP formulations for mucosal drug delivery.

To optimize nanoparticle formulations for mucosal delivery, several assays have been established to directly measure mucus penetration efficiency. Early work primarily used diffusion chambers where mucus collected from animals or humans is placed between donor and acceptor compartments where the fraction of particles that reach the acceptor chamber is monitored over time.^24^ Microscopy–based methods such as fluorescence recovery after photobleaching (FRAP) and particle tracking are now often used to directly measure nanoparticle diffusion within mucus.^25–28^ However, the methods available to assess NP transport through the mucus barrier are generally low-throughput which limits the ability to directly compare a wide range of formulation strategies. Moreover, it is often difficult to acquire mucus samples from humans to perform these assessments. Given the wide range of design parameters, it would be desirable to assess many NP formulations in parallel to down-select potential mucosal delivery strategies for further evaluation. In addition, the high-throughput system should be able to capture the changes in mucus properties as a function of tissue type and disease state.

Towards this end, we report a new strategy to screen nanoparticle formulations for mucosal delivery applications. Specifically, we developed a synthetic mucus barrier array (SMBA) platform containing mucin-based hydrogels which can be tailored to mimic the viscoelastic properties of native mucus in health and disease.^29–31^ To confirm the validity of the SMBA system, we performed studies using polystyrene (PS) NP with size and surface chemistries previously evaluated in the literature. Based on the use of NAC as a mucolytic agent to enhance the penetration of NP through hyper-concentrated mucus produced in cystic fibrosis lung disease,^16,32^ we then evaluated a previously untested mucolytic agent TCEP to determine if it could potentially serve as an NP permeation enhancer. Our data establishes proof-of-concept SMBA can be used to screen candidate NP formulations prior to further *in vitro* and *in vivo* evaluation for oral, inhaled, and topical drug delivery applications.

## MATERIALS AND METHODS

### Synthetic mucus hydrogel formulation

The synthetic mucus (SM) hydrogel used within the arrays was previously developed to mimic the material properties of human mucus.^29^ A solution of 4% porcine gastric mucins (PGM; Sigma Aldrich; mucin from porcine stomach, type III, bound sialic acid 0.5-1.5%, partially purified powder) was stirred for 2 hours in a physiological buffer representative of the ionic concentrations found in mucus (154 mM NaCl, 3 mM CaCl_2_, and 15 mM NaH_2_PO_4_ at pH 7.4). Four arm-PEG-thiol (PEG-4SH, 10 kDa; Laysan Bio) was used as a crosslinking agent to form disulfide bonds between the mucins. A 4% solution of PEG-4SH was prepared separately using the same physiological buffer and combined in an equal volume ratio with the 4% PGM solution. The resulting SM hydrogels consisted of 2% PGM and 2% PEG-4SH, and are referred to as 2% SM gels. In experiments that utilized a 4% SM gel to mimic disease-state mucus, 8% PGM and 8% PEG-4SH solutions were prepared in the same manner and combined in equal volume ratios.

### Synthetic mucus barrier array design and preparation

Synthetic mucus barrier arrays (SMBA) were designed using computer-aided design software (Fusion360) with 9 wells that fit into an underlying 96-well flat black plate (Costar). The SMBA devices were 3D printed using an SLA Formlabs 2 printer with V4 white resin material. The SM hydrogels were cast so they achieved gelation within the wells of the array. In order to cast hydrogels into the SMBA, parafilm was stretched across the surface of the 96-well plate, then the bottom of the wells of the SMBA were pressed downward into the wells of the plate. This technique formed a tight seal of parafilm over the bottom of the SMBA wells. The hydrogel solution was added to the bottom of each well of the SMBA in the 96-well plate to allow for gelation in an upright position with the parafilm kept taut. The standard volume of hydrogel solution added to each well of the array devices was 30 μl, unless otherwise indicated. Assuming the gel maintains a cylindrical geometry within the SMBA, the hydrogel layer would possess a thickness of ∼2 mm. The 96-well plate containing the SMBA was incubated for 22 hours in a humidified chamber to prevent drying out of the hydrogels during gelation. The SMBA was then taken out of the 96-well plate with care taken to ensure that the gels were not disturbed while peeling them off the parafilm. The standard volume of 30 µl was found to be the minimum volume that could be used where the gels would remain intact in the SMBA for the duration of experiments (up to 2 hours). Use of volumes less than <30 µl lead to incomplete gel coverage and leakiness within the SMBA which precluded testing at shorter total mucus gel depths.

### Nanoparticle preparation

NPs were rendered muco-inert with a dense surface coating of polyethylene glycol (PEG). Carboxylate modified fluorescent PS NPs with diameters of 20 nm, 100 nm, and 500 nm (Life Technologies) were coated with PEG. Five kDa methoxy PEG-amine (Creative PEGworks) was attached to the surface of the NPs by a carboxyl-amine linkage, as previously described,^28,29^ The zeta potential of PEG-coated NPs (PS-PEG NPs) was measured using a Nanobrook Omni Particle Analyzer (Brookhaven Instruments). NPs were considered adequately PEGylated if measured zeta potential was near neutral in charge.

### Determining nanoparticle penetration efficiency using SMBA

SMBAs with solidified gels were placed into a fresh 96-well black plate (Costar) with 100 μl of PBS added to the wells of the plate. NPs were diluted in PBS prior to addition to the SMBA. The standard volume of NP solution added to the SMBA wells, unless otherwise indicated, was 10 μl. In experiments in which the mucolytics TCEP or NAC are used, the mucolytic was added directly into the solution of NPs in PBS at a concentration of 10 mM. As shown in **Figure 1**, the gels were submerged in the 96-well plate containing PBS. The gels were then incubated after the addition of the NP solution for 2 hours at room temperature before the SMBAs were removed from the 96 well plate. Positive control wells were also included in the 96-well plate, where the same NP solution added to the SMBA wells was added directly to the PBS in the 96-well plate. The positive control wells were used as a reference to determine the fluorescence of the full amount of NPs added to each SMBA well, simulating 100% penetration of NPs, to aid in calculating the percent penetration of the NPs across the gels in the experimental wells of SMBAs. A standard curve of serial dilutions of PS NPs from the stock solution was made in the same 96-well plate to allow for conversion between fluorescence units and known NP concentration. Then, a fluorescence reading was taken to assess the concentration of NPs in the PBS within the 96-well plate. The percentage of NPs to penetrate the gels in the experimental wells was calculated by converting the fluorescence units measured in each well to NP concentration based on the standard curve. Finally, the percent penetration of NPs across the gels was found by dividing the concentration in the experimental wells by the averaged concentration of NPs in the positive control wells.

**Figure 1.**
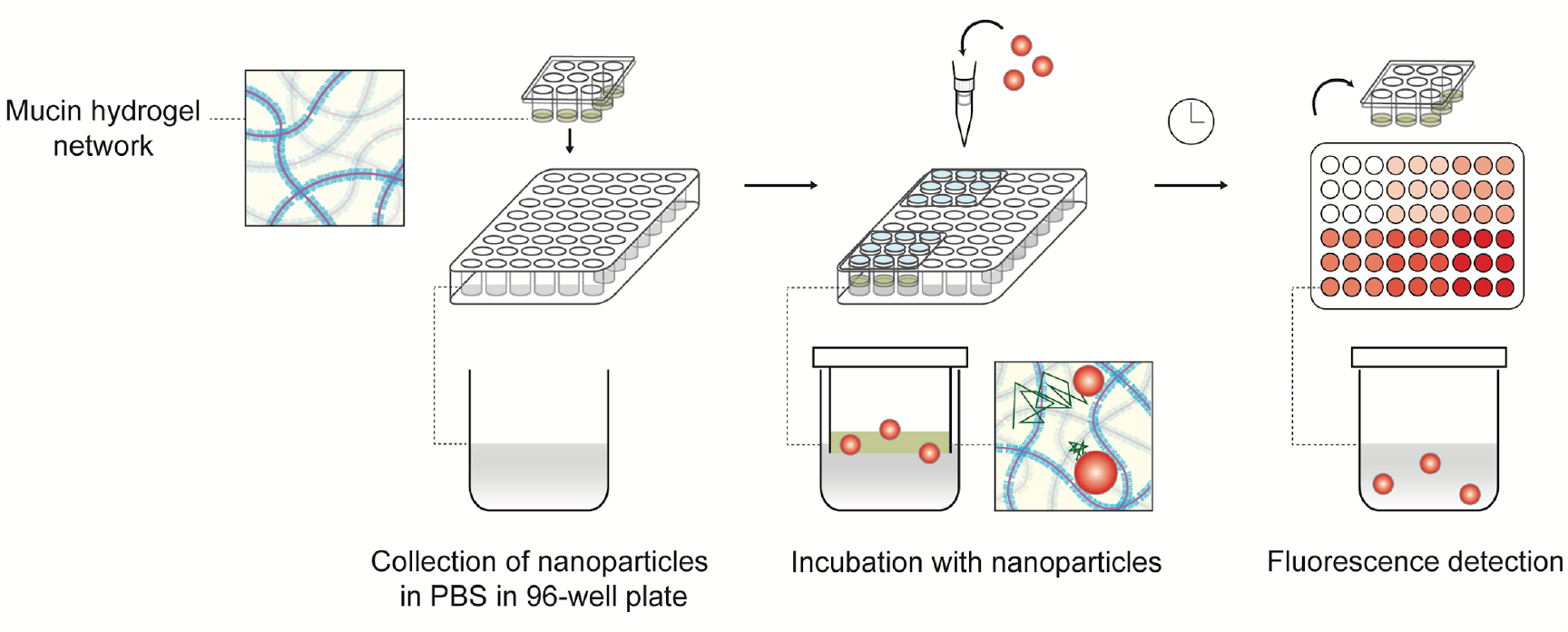
Schematic of SMBA experimental setup. SM hydrogels are cast in the wells of the SMBA device, then placed in a 96 well plate containing PBS. A solution of fluorescent NPs is added to the apical surface of the gels. NPs that penetrate the SM gel are detected via fluorescence.

## Results & Discussion

### NP size and surface chemistry affect their penetration through SMBA

For initial validation of the SMBA screening system, several parameters known to impact nanoparticle penetration through the mucus barrier were assessed (**Figure 2A)**. As noted, NP charge and hydrophobicity must be optimized to avoid adhesive interactions with the mucus barrier.^10,11^ To assess the effect that NP surface chemistry has on penetration, 20 nm and 100 nm NPs were coated with poly-ethylene glycol (PEG) to neutralize their surface charge. Carboxylate-modified PS and PS-PEG NPs were added topically to the 2% SM hydrogels within the SMBA system and the fraction of NP to cross the synthetic mucus barrier in 2 hours was measured (**Figure 2B)**. PEGylated NPs in both sizes (20 nm and 100 nm) displayed significantly higher percent penetration than their non-PEGylated counterparts in the SMBA system. We then evaluated the effect of NP size on particle penetration. Percent penetration of 20 nm, 100 nm, and 500 nm PS-PEG NPs across the SMBA system was compared (**Figure 2C)**. The results seen were consistent with prior studies^17,23,28^ as we observed a negative correlation between nanoparticle size and particle penetration across the synthetic mucus barrier. These experiments validate that the SMBA system can accurately predict how these conditions impact particle penetration, by demonstrating that changes in NP surface chemistry and size will affect particle penetration in a manner consistent with previous studies.

**Figure 2.**
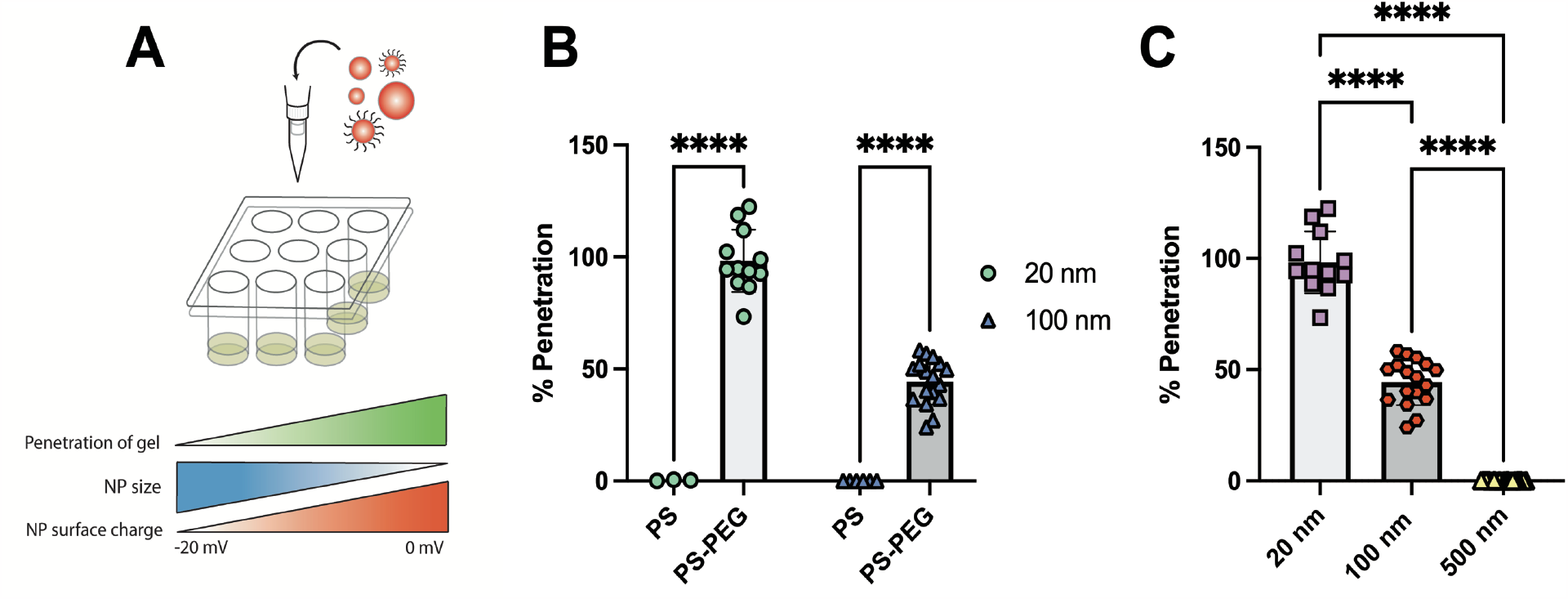
NP size and surface chemistry affect their penetration through SMBA. A) Schematic of the effects of NP size and surface charge on SM gel penetration. B) Comparison the percentage of PS NPs (highly negative surface charge) versus PS-PEG NPs (neutral surface charge) to penetrate 2% SM gels in the SMBA device, repeated with both 20 nm and 100 nm NPs. C) Penetration of PS-PEG NPs across the gels in the SMBA device with various diameters (20 nm, 100 nm, 500 nm).

### Impact of barrier thickness and solution volume on NP penetration through SMBA

To further assess NP penetration through SMBA under varying conditions, we next determined the effect of gel thickness and NP solution volume on NP penetration (**Figure 3A)**. To vary the gel thickness, 2% SM gels were cast in the SMBA device in two different volumes of either 30 μl, designated as a low gel volume, or 50 μl, designated as a high gel volume. Both 20 nm and 100 nm PS-PEG NPs were then added topically to the SM hydrogels within the SMBA system and percent particle penetration was evaluated (**Figure 3B)**. A negative correlation was seen between the SM hydrogel volume and NP penetration of the gel as both 20 nm and 100 nm PS-PEG NP groups displayed significantly lower percent particle penetration in the higher gel volume (50 μl) SM hydrogels within the SMBA system. This can likely be attributed to a larger effective distance that the NP must travel to penetrate a mucus barrier of greater thickness. We then analyzed the effect of NP solution volume administered topically to the SM hydrogels within the SMBA system **(Figure 3C)**. We examined percent particle penetration across 2% SM gels within the SMBA system of both 20 nm and 100 nm PS-PEG NPs suspended in PBS solution in dosage volumes of either 10 μl, a relatively low NP volume, or 20 μl, a relatively high NP volume. A positive correlation was seen between the size of the NP solution administered and the percent of particle penetration seen in the 20 nm NP group, as a high NP solution volume (20 μl) had a statistically significantly higher percentage of NP penetration when compared to the low NP solution volume (10 μl). It is important to note that although this trend was seen, both groups had a high percent particle penetration (∼90-100%). Although a similar trend was observed within the 100 nm NP group, no statistically significant difference was observed between the high NP solution volume and low NP solution volume groups. Thus, changing the volume of NP administered should not significantly impact the ability of the SMBA to detect differences in NP penetration between different conditions.

**Figure 3.**
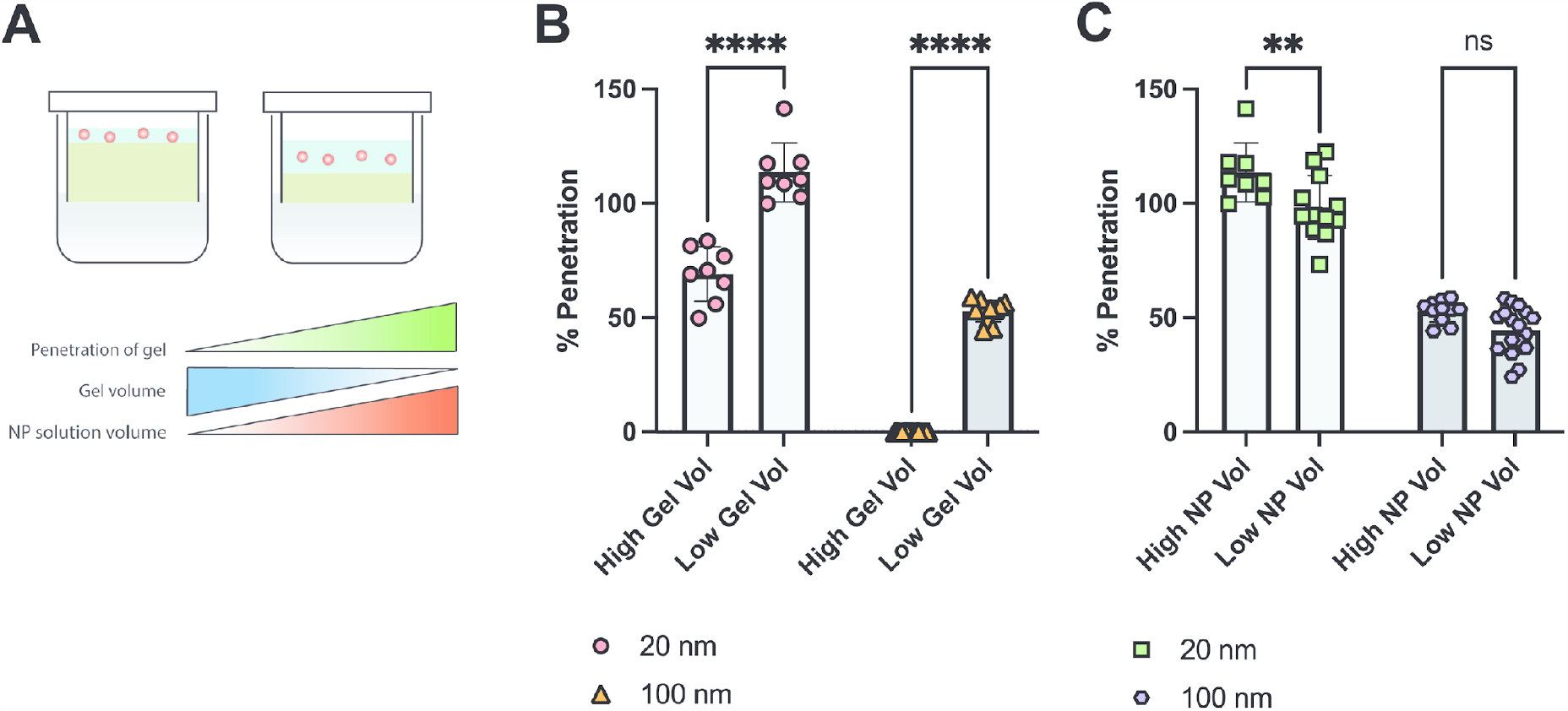
Impact of barrier thickness and solution volume on NP penetration through SMBA. A) Schematic of the effects of varying either the SM gel volume or NP solution volume added to the SMBA device on diffusion across the SM gels. B) 2% SM gels were cast in the SMBA device in two different volumes of either 30 l (low gel vol) or 50 l (high gel vol), then the percentage of both 20 nm and 100 nm PS-PEG NPs to penetrate the gels was quantified. C) Percent penetration across SM gels in the SMBA device of both 100 nm and 20 nm PS-PEG NPs suspended in PBS solution applied in two different volumes of either 10 µl (low NP vol) or 20 µl (high NP vol). Statistical significance determined by two-way ANOVA with Šídák’s multiple comparisons test (B,C). (ns = p > 0.05, **** = p < 0.0001, ** = p < 0.01).

### Evaluating mucolytics as permeation enhancers for inhaled NP delivery using SMBA

In many chronic lung diseases such as asthma and cystic fibrosis, hyper-concentrated mucus is produced which may further limit the penetration of NP delivery systems under evaluation for inhaled drug and gene delivery.^25,34,35^ To model disease-like conditions, we increased the total gel solids concentration and then assessed NP penetration using SMBA (**Figure 4A)**. We prepared 2% w/v synthetic mucus gels to represent airway mucus in health and 4% w/v synthetic mucus gels to represent mucus in individuals with obstructive lung disease.^23,36,37^ We observed that percent particle penetration of the 100 nm PS-PEG NPs was significantly reduced in the 4% w/v SMBA when compared to the healthy state 2% w/v SMBA (**Figure 4B)**. These results highlight the potential utility of SMBA to examine the impact of alterations to the mucus barrier in disease. Given the significantly limited penetration of 100 nm NP under disease conditions, we then tested the impact of reducing agents, often used as mucolytic therapies, to enhance NP permeation through SMBA. We compared two mucolytic agents used as therapeutics to improve clearance of airway mucus: N-acetylcysteine (NAC), and Tris (2-carboxyethyl) phosphine (TCEP).^34,38,39^ Both NAC and TCEP act as mucolytics by reducing mucin-mucin disulfide bonds which directly degrades the mucus gel and reduces its viscoelasticity. NAC has been previously used in conjunction with PEGylated NP where it has been shown to enhance NP penetration through the airway mucus barrier.^16,32^ However to our knowledge, TCEP has yet to be tested as a permeation enhancer to improve inhaled NP delivery. We hypothesized TCEP would significantly enhance NP penetration in comparison to NAC as it has been shown to possess a much higher activity at reducing mucin biopolymers.^38,40^ To test this, we formed 4% w/v SMBA (disease state) and treated them with either NAC or TCEP and visually observed a significant degradation of the disulfide-linked synthetic mucus gel **(Figure 4C)**. We then examined the percent of particle penetration of 100 nm PS and PS-PEG NPs when applied to 4% SM gels in the SMBA system in combination with either 10 mM TCEP or 10 mM NAC **(Figure 4D)**. We observed that both PS and PS-PEG NPs can achieve significantly greater penetration in SM hydrogels in which TCEP is present when compared to SM hydrogels in which NAC is present. The results of these studies suggest TCEP may be a better alternative to NAC given the observed improvements in NP penetration through the mucus barrier. Studies can also be conducted in the future to further optimize the concentration of TCEP required to enhance NP penetration through the mucus barrier.

**Figure 4.**
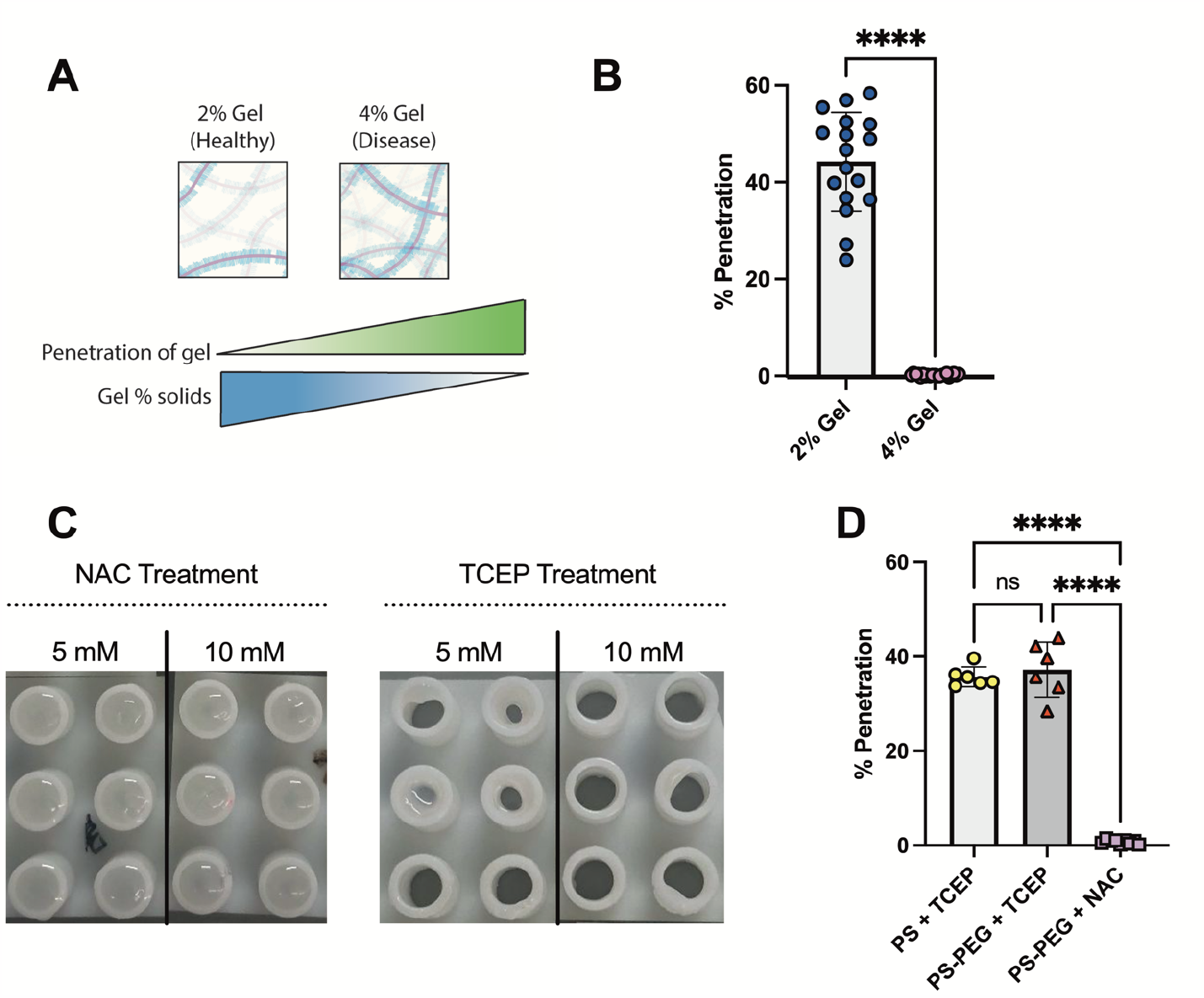
Evaluating mucolytics as permeation enhancers for inhaled NP delivery using SMBA. A) Schematic of the effects of gel solids on NP penetration of the SM gel. B) Comparison of 100 nm PS-PEG NP penetration through either 2% SM or 4% SM gels in the SMBA devices representing healthy and disease state mucus, respectively. C) Images of the bottom of the SMBA devices containing 4% SM gels after treatment with the mucolytics NAC or TCEP at both 5mM and 10 mM concentrations. D) Percent penetration of 100 nm PS or PS-PEG NPs when applied to 4% SM gels in the SMBA device in combination with either 10 mM TCEP or NAC. Statistical significance determined by unpaired t-test (B) and one-way ANOVA with Tukey’s multiple comparisons test (D). (ns = p > 0.05, **** = p < 0.0001).

## Conclusions

We demonstrated that the SMBA system can be used to predict the fate of nanomedicine in the mucus barrier. By examining the effects of NP surface chemistry and size as well as mucus barrier concentration and thickness, we were able to demonstrate the utility of the SMBA to evaluate nanoparticle-based therapeutics targeted toward mucosal environments. Our head-to-head comparison of NAC and TCEP as NP permeation enhancers highlighted how SMBA could be helpful in optimizing formulation strategies by considering disease-associated changes to the mucus barrier. This proof-of-concept study will be expanded in future research to other clinically relevant drug and gene delivery systems (e.g. biodegradable polymeric NP, lipid NP, extracellular vesicles, viral vectors)^41–45^ to optimize their properties for mucus penetration and improved therapeutic effectiveness. Ultimately, this work provides a simple but powerful method to assess NP design strategies for therapeutic applications targeting mucosal tissues.

## Declaration of Competing Interest

The authors declare no conflict of interest.

## Acknowledgements

This work was supported by the Burroughs Wellcome Fund Career Award at the Scientific Interface (to G.A.D.), Cystic Fibrosis Foundation (BOBOLT23H0 to A.B.), the National Science Foundation (2047794, 2129624 to G.A.D.), and the National Institutes of Health (R21AI142050 to G.A.D.).

## Notes

### Competing Interest Statement

The authors have declared no competing interest.

